# Effect of user adaptation on prosthetic finger control with an intuitive myoelectric decoder

**DOI:** 10.1101/585703

**Authors:** Agamemnon Krasoulis, Sethu Vijayakumar, Kianoush Nazarpour

## Abstract

Machine learning-based myoelectric control is regarded as an intuitive paradigm, because of the mapping it creates between muscle co-activation patterns and prosthesis movements that aims to simulate the physiological pathways found in the human arm. Despite that, there has been evidence that closed-loop interaction with a classification-based interface results in user adaptation, which leads to performance improvement with experience. Recently, there has been a focus shift towards continuous prosthesis control, yet little is known about whether and how user adaptation affects myoelectric control performance in dexterous, intuitive tasks. We investigate the effect of short-term adaptation with independent finger position control by conducting real-time experiments with 10 able-bodied and two transradial amputee subjects. We demonstrate that despite using an intuitive decoder, experience leads to significant improvements in performance. We argue that this is due to the lack of an utterly natural control scheme, which is mainly caused by differences in the anatomy of human and artificial hands, movement intent decoding inaccuracies, and lack of proprioception. Finally, we extend previous work in classification-based and wrist continuous control by verifying that offline analyses cannot reliably predict real-time performance, thereby reiterating the importance of validating myoelectric control algorithms with real-time experiments.

## Introduction

State-of-the-art commercial prosthetic hands exhibit hardware capabilities that could potentially allow their users to independently control individual fingers. However, this feature is almost never utilised; instead, most current prosthetic systems still employ the conventional amplitude-based, dual-site electromyogram (EMG) mode switching paradigm for grip selection and actuation^1^. Due to using a highly-non-intuitive control interface, the efficacy of this method relies on user experience gathered during daily interaction with the device. It has been previously shown, and is currently well-accepted, that humans are capable of greatly improving their control of mode switching myoelectric interfaces within only a few days of training^2,3^.

Moving towards more natural interfaces, a large body of research has investigated the potential of pattern recognition methods for providing users with the ability to directly access desired control modes, such as hand grips and/or wrist functions. The idea of EMG signal classification for prosthetic control has been around for almost half a century4 and has recently found its way into commercial adoption^1^. Although classification-based control is regarded as an intuitive scheme, there has been evidence that experience can still lead to substantial performance improvement^5–8^. This can be attributed to various causes, for example, an increase in class separability5 and movement repeatability^6^, even in the absence of any form of feedback^7^. Recently, a clinical study involving transhumeral amputees having undergone targeted muscle reinnervation reported a significant increase in classification-based myoelectric control performance within two months of daily use^8^.

The grip classification approach can offer a remarkable improvement of the intuitiveness and ease of use of the prosthetic device, however, it suffers from two main limitations: 1) it results in severe under-actuation of the prosthesis, which dramatically limits its functionality, as the user can only have access to a set of pre-determined modules; and 2) it is sequential in nature, that is, a single class of movement can be active at a time, as opposed to the natural continuous and asynchronous finger movement exhibited by the human hand. One way of enhancing the dexterity of powered myoelectric prostheses is via continuous and simultaneous control of multiple degrees of freedom (DOFs)^9^. Arguably, the primary focus of continuous myoelectric control has previously been on restoring wrist function^10–13^. Nevertheless, over the last decade several groups have also addressed the challenge of using surface EMG signals to reconstruct kinematic variables (e.g. position or velocity) of independent finger movement, both offline^14–18^ and in real-time^19–21^. As compared to non-invasive methods, intramuscular recordings offer the advantage of lower level of muscle cross-talk^22^, hence making it possible to create multiple one-to-one mappings between specific muscles and prosthesis degrees of actuation (DOAs). This opportunity has been explored in the context of controlling both virtual23 as well as prosthetic24 hands. Besides finger position and velocity decoding, individual fingertip forces have also been reconstructed offline^25,26^ and in real-time^27–29^ using surface EMG signals.

Continuous myoelectric control strategies, which are often referred to as *proportional*^9^, are typically intuitive, that is, they operate based on physiological associations between muscle (co)-activation patterns and prosthesis DOAs. They usually require an initial phase of data collection for subject-specific model training. A promising alternative is based on user adaptation^30^, whereby muscle signals control the prosthesis DOAs using pre-defined, subject-independent mappings. This approach heavily relies on user adaptation taking place during closed-loop control, therefore the provision of continuous feedback (e.g. visual) during user training is necessary. There has been increasing evidence that humans are able to develop novel task-specific muscle synergies, that is, muscle co-activation patterns, to achieve high-level performance in a variety of tasks, including two-dimensional cursor position control^30–35^, prosthetic finger position^32^, and high-dimensional robotic arm control^36,37^. Notably, it has been found that such synergistic patterns can be learnt even when they are not intuitive from a physiological perspective, for instance, due to requiring the co-activation of antagonist muscles^31^.

In comparison with non-intuitive interfaces, whereby an inverse model has to be learnt from scratch, the effect of user experience on myoelectric control performance when using an intuitive, regression-based approach is much less understood. In the context of 2-DOF continuous wrist control, a previous study showed that while three machine learning algorithms yielded statistically different offline decoding accuracies, the performance of the three algorithms was comparable during real-time myoelectric control^12^. Additionally, only weak, mainly non-significant correlations were observed between offline and real-time control performance measures. These findings support the view that user adaptation mechanisms that take place during closed-loop interaction affect ultimate real-time control performance, thereby questioning the extent to which offline myoelectric control studies can inform clinical translation of advanced upper-limb prostheses. Similar findings have been also reported in the context of myoelectric classification^38,39^. Furthermore, a study showed that real-time, regression-based prosthetic wrist control might be less susceptible to perturbations, for example, due to noise in EMG signals, than its offline decoding counterpart^40^. This observation provides evidence that humans can user error correction mechanisms to compensate for decoding inaccuracies during closed-loop interaction with myoelectric interfaces. With regard to prosthetic finger control, several studies have attempted to push the boundaries of offline decoding accuracy^16,18^. However, substantially less effort has been made towards understanding whether and in what manner user adaptation can affect real-time control performance.

In this work, we investigate the effect of user adaptation in continuous prosthetic finger control in able-bodied and transradial amputee subjects. We hypothesise that however intuitive a myoelectric task might be, experience gathered during interaction with the interface would still lead to performance improvement. We evaluate our hypothesis using two intuitive finger control schemes, namely, EMG-based finger position control and teleoperation with an instrumented data glove. Additionally, we investigate the effect of user experience on the power of the recorded EMG signals and the variability of the controllable DOAs. Finally, we extend previous work on myoelectric classification and continuous wrist control^38,39^, by demonstrating that it is not possible to reliably predict real-time prosthetic finger control performance solely based on the outcomes of offline decoding analyses. To the best of the authors’ knowledge, this is the first study to systematically demonstrate the positive impact of short-term adaptation, achieved through biofeedback user training, on intuitive, dexterous prosthetic finger control both with EMG-based decoding, as well as during robotic hand teleoperation with a data glove.

## Materials and Methods

### Participant recruitment

Ten able-bodied (nine male, one female; all right-hand dominant; median age, 26.5 years) and two male, right-hand transradial amputee subjects were recruited. Both amputees were right-hand dominant prior to amputation. Some of the able-bodied (five out of ten) and both amputee participants had previously taken part in classification-based myoelectric control experiments^41^.

### Ethics statement

All experiments were performed in accordance with the Declaration of Helsinki and were approved by the local Ethics Committees of the School of Informatics, University of Edinburgh and School of Engineering, Newcastle University. Prior to the experiments, subjects read a participant information sheet and gave written informed consent.

### Hardware and signal acquisition

For the able-bodied group, 16 Delsys Trigno^TM^ sensors (Delsys, Inc.) were placed on the participants’ right forearm arranged in two rows of eight equally spaced sensors, without targeting specific muscles. The two rows were placed 3 cm and 5.5 cm, respectively, below the elbow. Photographs showing electrode placement for an able-bodied participant are shown in Figure 1A,B. Using a similar configuration, 13 and 12 sensors were used, respectively, for the two amputee participants, due to limited space availability on their remnant limb (right limb in both subjects). Prior to sensor placement, participants’ skin was cleansed using 70% isopropyl alcohol. Elastic bandage was used to secure the sensor positions throughout the experimental sessions. Following sensor placement, the quality of all EMG channels was verified by visual inspection. The sampling frequency of EMG signals was set to 1111 Hz.

**Figure 1.**
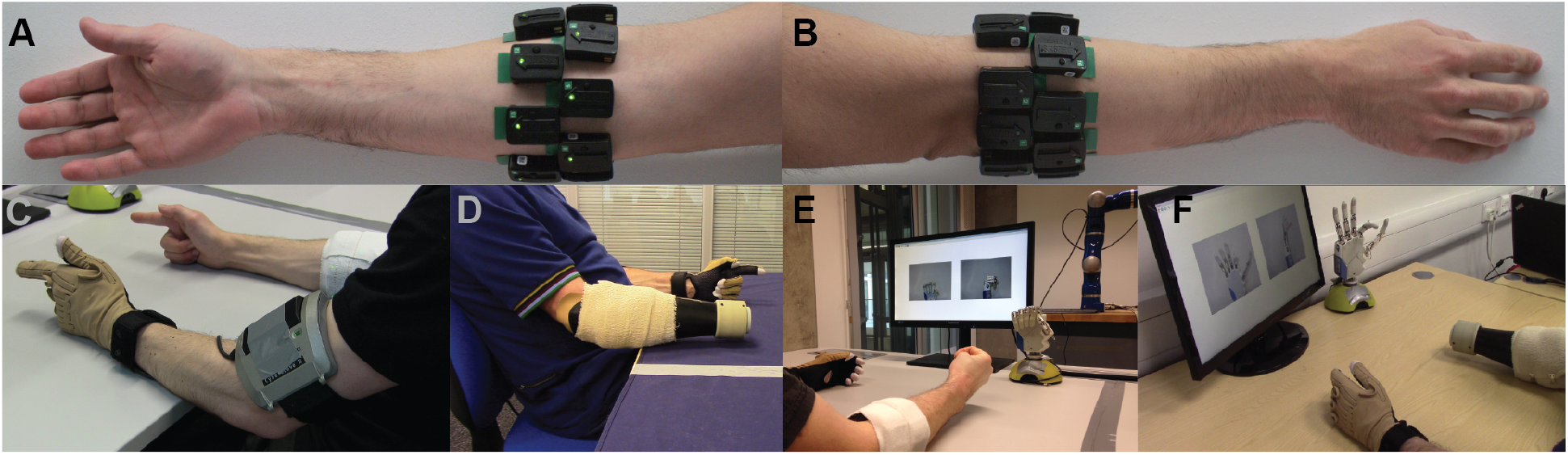
Experimental setup. Surface EMG electrodes were placed on subjects’ forearm below the elbow in two rows of equally spaced electrodes. (**A**) Palmar and (**B**) dorsal views of the forearm. Refer to main text for details on number of electrodes used for able-bodied and amputee participants. (**C**)-(**D**) Bilateral mirrored movement training. Able-bodied and amputee participants shown during initial data collection. Muscle activity was recorded from the participants’ right forearm (i.e. the remnant limb for amputees), whereas hand kinematic data were recorded from the participants’ left hand with an 18-DOF data glove. (**E**)-(**F**) Real-time posture matching task. Able-bodied and amputee participants shown while they modulate their muscle activity to control the finger positions of the robotic hand. The target postures for the shown trials were (**E**) full cylindrical grip and (**F**) half index flexion.

An 18-DOF CyberGlove II data glove (CyberGlove Systems LLC) was used to record hand kinematic data from the participants’ left hand (Figure 1C,D). For each participant, the glove was calibrated prior to data collection using dedicated software provided by the manufacturer. The sampling rate of glove data was set to 100 Hz.

For the real-time control experiments, we used a right model of the IH2 Azzurra hand (Prensilia s.r.l.), which is an externally-powered, underactuated (11 DOFs, 5 DOAs) anthropomorphic hand. It comprises 4 intrinsic motors controlling flexion and extension of the five digits (the ring and little fingers are mechanically coupled) and an additional motor controlling thumb rotation. The robotic hand is shown in Figure 1E,F.

### Experimental design

Participants sat comfortably on an office chair and rested both arms on a computer desk. Each participant completed one experimental session, which comprised two main phases: 1) *initial data collection,* and 2) *real-time robotic hand control.* Each experimental session lasted around 140 minutes, which included: skin preparation, electrode positioning, and signal inspection (20 minutes); initial data collection (60 minutes); short interval (20 minutes); and real-time control of the robotic hand (40 minutes).

### Initial data collection

In the first part of the experiment, participants were asked to reproduce a series of motions instructed to them on a computer monitor. Nine exercises were selected for data collection ranging from individuated-finger to full-hand motions. The nine motions comprised: thumb flexion/extension, thumb abduction/adduction, index flexion/extension, middle flexion/extension, ring/little flexion/extension, index pointer, cylindrical grip, lateral grip, and tripod grip (Supplementary Figure S1). All participants were asked to perform bilateral mirrored movements with both their arms resting on a computer desk.

We recorded three datasets (i.e. separate blocks of trials) for each participant during this phase of the experiment: the first two *(training* and *validation* sets) comprised 10 repetitions of each motion, and the third one *(testing* set), only two. Each motion execution lasted approximately 7 s and at the end of each trial subjects were instructed to return to the rest pose which corresponded to complete muscle relaxation (shown in Supplementary Figure S1A). Succeeding trials were interleaved with intervals of 3 s and participants were also given a 10-minute break after the completion of each block of trials.

### Signal pre-processing

We used a sliding window approach to process the EMG signals. The length of the window was set to 128 ms with an increment of 50 ms (60% overlap). The following time-domain features were extracted from the recorded EMG channels: Wilson amplitude, 4^th^-order auto-regressive coefficients, waveform length, log-variance, and slope sign change^11,42^. The columns of the design (i.e. feature) matrices were subsequently standardised via mean subtraction and inverse standard deviation scaling. Feature means and standard deviations were estimated using training data only.

For the hand kinematic data that were recorded with the data glove, we computed the mean value within the processing window for each DOF individually. The calibrated glove measurements were converted into digit positions for the prosthetic hand using a linear mapping (see Supplementary Methods). The columns of the target matrices containing the prosthetic hand joint positions were finally normalised in the range [0,1], where *y_j_* = 0 corresponds to full extension and *y_j_* = 1 to full flexion of the *j*^th^ DOA, respectively.

### Model training, prediction post-processing, and hyper-parameter optimisation

Model training took place during the short resting interval between the initial data collection and real-time control evaluation. To decode finger positions from muscle activity, we deployed a regularised version of the Wiener filter, implemented using auto- and cross-correlation matrices^43^, which we have previously used to reconstruct finger position trajectories from myoelectric data offline^17,44^. We have shown in previous work that the generalisation of this decoder is comparable to that of non-linear regression algorithms when tested on movements outside the training set^17^. The Wiener filter is a classical signal processing method for estimating a target variable using linear time-invariant filtering (e.g. spatial or temporal). In other words, at time instance *n*, each input *x_d_* (i.e. EMG feature) is convolved with a finite impulse response function to produce an output y (i.e. digit position of a single DOA of the prosthetic hand):

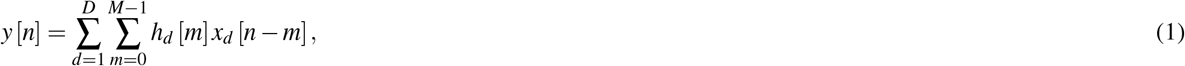

where *h_d_* [*m*] accounts for the contribution of the input *d* at time instance *m, x_d_* [*n – m*] is the activation of the input d at time *n – m, M* is the filter length, and we also assume a finite number of samples *n* = 1,…, *N*. We set the length of the linear filters to 300 ms, which corresponds to including at each time step *M* = 6 previous time lags, assuming a fixed window increment of 50 ms. The number of EMG electrodes used for decoding varied across subjects and was based on a sequential selection algorithm (described below). When the full set of sensors was used, the input dimensionality was *D* = 672 (i.e. 7 EMG features / (electrode × time bin) × 16 electrodes × 6 time bins). The output dimensionality was *K* = 5, that is, the number of DOAs of the robotic hand. For covariance matrix estimation, we used L_2_ regularisation to avoid inversion of potentially ill-conditioned matrices due to the high dimensionality of the input space.

We post-processed predictions using exponential smoothing to ensure smooth digit trajectories. We implemented this in the time-domain as follows:

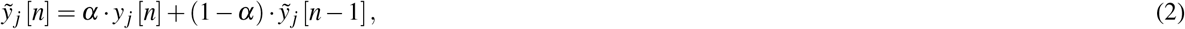

where *y_j_* [*n*] and 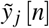 denote, respectively, the raw and smoothed predictions of the *j*^th^ DOA at time step *n*, and *α* is the smoothing parameter, which is constrained by 0 ≤ *α* ≤ 1. Smaller values of *α* result in stronger smoothing, that is, a smaller cutoff frequency when smoothing is viewed as a low-pass filter, but also increase the prediction response latency.

We performed three types of model selection (i.e. hyper-parameter tuning) for each participant during the training phase: sensor selection, regularisation, and smoothing parameter optimisation. Models were initially trained using data from the training set only. Model selection was carried out by means of maximising performance on the validation set. Following parameter optimisation, the training and validation sets were merged and used to train final models. The test set was only used to evaluate and report offline performance of the final models.

Offline reconstruction accuracy was assessed using the multivariate coefficient of determination (R^2^) metric defined as:

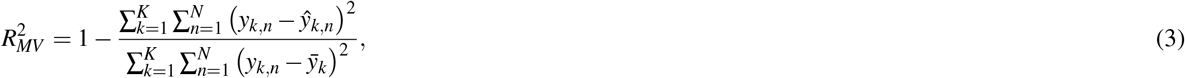

where *K* denotes the dimensionality of the target variable (in our case *K* = 5), *N* is the number of samples in the measurement/prediction vector, *y_k,n_* and *ŷ_k,n_* are the *n*^th^ observed and predicted values of the *k*^th^ output variable, respectively, and 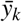 denotes the sample mean of the *k*^th^ output variable.

For sensor selection, a standard sequential forward method was used^45^. That is, the selection algorithm started with an empty set and at each iteration the sensor that yielded the highest reconstruction accuracy improvement was added to the pool. The algorithm terminated execution when the inclusion of any remaining sensors caused a decrease in average performance. The number of used sensors varied from 8 to 16, with a median value of 12. To optimise the regularisation parameter λ of the Wiener filter, a search was performed in the log-space {10^-6^, 10^-5^,…, 10^1^} using a factor (i.e., multiplicative step) of 10. Finally, the exponential smoothing parameter *α* (Equation (2)) was optimised via linear search in the range [0,…, 1] with a step size of 0.01. The three model selection steps were performed sequentially in the following order: sensor selection, λ optimisation, and *α* optimisation. In other words, the subset of sensors was firstly identified; using the selected subset, the regularisation parameter λ was tuned; finally, using the selected sensor subset and chosen value for λ, the smoothing parameter *α* was optimised.

### Real-time control and evaluation

A real-time, biofeedback, posture matching task was designed to assess the efficacy of the regression-based control scheme and provide insight into the effect of user practice on prosthetic finger control. To that end, two scenarios were investigated: in *EMG control mode,* participants were required to modulate their muscle activity to control the prosthetic hand by making use of the regression model; in *glove control mode,* participants teleoperated the hand using the data glove worn on their contralateral hand, that is, the intact limb for the amputee participants. The glove control mode was included for two reasons: to provide an estimate of the upper-bound of prosthetic control performance for the designed experimental task (i.e., *benchmark);* and to investigate whether user practice leads to performance improvement during prosthetic finger control with a direct, natural control interface.

Participants were presented with a series of target postures on the screen and were instructed to control the hand to match the desired postures as closely as possible. Image prompts were only used at this stage, as opposed to the training data collection phase, where participants were instructed to follow video prompts. During the task, the robotic hand was connected to a base stand placed on the surface of the desk and sitting in front of the participant (Figure 1E,F). Nine hand postures were included, each of them with two variations, half and full activation. Therefore the total number of postures in the real-time experiment was 18. The included hand postures were: thumb abduction, thumb flexion, index flexion, middle flexion, ring/little flexion, index pointer, cylindrical grip, lateral grip, and pinch grip (Supplementary Figure S2).

At the start of each trial, the participants were presented with a pair of static pictures providing front and side views of the desired posture. An audio cue (waveform, sine wave; frequency, 400 Hz; duration, 500 ms) was used to signal the initiation of the trial. Participants were then given 3.5 s to drive the prosthetic hand into the desired posture. At the end of this period, a second audio cue (waveform, sine wave; frequency, 800 Hz; duration, 500 ms) was used to signal the initiation of the evaluation phase of the trial, which lasted 1.5 s. During the evaluation phase, participants were instructed to hold the hand in the performed posture. At the end of the evaluation phase, the hand was reset to its initial posture (i.e., fully open) signalling the end of the trial. Pictures illustrating the real-time posture matching task are shown in Figure 1E,F for two participants, one able-bodied and one amputee.

At the end of each trial, participants received a score characterising their performance. This score was based on the average mean absolute error (MAE) between the target and performed postures during the evaluation phase (i.e. the last 1.5 s) of the trial. Let ***y*** and ***ŷ*** denote K-dimensional vectors in a real vector space. In our case, the two vectors represent the target and performed postures, respectively, of the prosthetic hand at a given time step and *K* = 5 is the number of DOAs of the hand. The MAE is defined as:

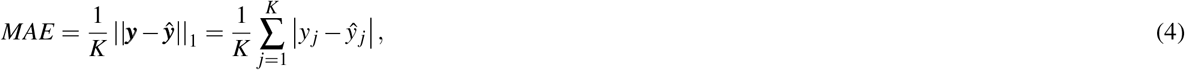

where *y_i_* and *ŷ_i_* denote, respectively, the target and true positions of the *j*^th^ DOA. The evaluation phase lasted for 1.5 s, and a finger position update was made every 50 ms, that is, the increment time of the sliding window. Thus, there were *N* = 300 distance samples associated with each trial. The average distance during the evaluation phase of a trial was estimated by computing the median across the samples of the population.

To provide the participants with an intuitive performance measure for each trial, MAEs were transformed into scores in the range of 0% to 100%. This transformation was achieved as follows: firstly, a baseline average MAE score between the target posture and random predictions was established by simulating 10^6^ random predictions uniformly sampled in the range [0,1]; the normalised score was then computed as:

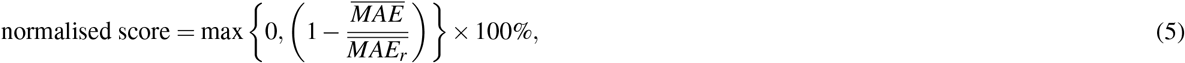

where 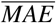 denotes the population average (i.e. median) MAE during the evaluation phase, and 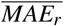 is the pre-computed, average random prediction error for the specified posture. This transformation ensured that a perfect reproduction of the desired posture would correspond to a 100% score, whereas a randomly performed posture would yield a score close to 0%. Negative scores were not allowed by the max operation. The random seed was controlled during the experiments to ensure the use of identical random predictions for all participants.

The posture matching task was split into several blocks. Within each block, all 18 postures were presented to the participants exactly once in a pseudo-randomised order. Each participant performed six blocks for each control mode (i.e. EMG and glove control, see Results section) therefore the total number of trials for each participant was 108 (i.e. 6 blocks × 18 trials/block). The execution of each block lasted approximately 3 minutes. At the end of block 3, participants were given a 1-minute rest. The stimulus presentation sequence was the same for all participants, but the order of the two control modes was counter-balanced across the two participant groups (i.e. able-bodied and amputee).

### Dimensionality reduction analysis

Dimensionality reduction analysis was performed by using principal component analysis (PCA) on the envelopes of the EMG signals. EMG envelopes were computed by using a sliding window approach (length 128 ms; window increment 50 ms) and extracting the mean absolute value within the window. All EMG electrodes were used in the analysis, regardless of whether they were used for decoding, therefore the dimensionality of the problem was equal to the total number of electrodes used for each participant (i.e. 16 for able-bodied participants, 13 and 12 for the first and second amputee subjects, respectively). To compare principal components (PCs) extracted from different experimental blocks, the absolute value of the cosine similarity was used, since the sign of PC directions was of no interest; in other words, two identical PC vectors with opposite signs were considered equivalent. The cosine similarity between two vectors ***a*** and ***b*** is defined as:

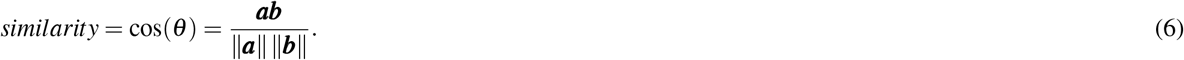

To compare PCs between two different blocks, the PCs of the blocks were first matched in terms of highest cosine similarity. The dimensionality reduction analysis was performed in Python using the scikit-learn library (v.0.19.1^)46^.

### Statistical analysis

For each participant, single trials were pooled together and used to compute subject-specific summaries. Depending upon the outcomes of D’Agostino-Pearson normality tests, the subject summaries used were either population means, for groups with samples following normal distributions, or medians, otherwise. The size of the summary groups equalled the total number of participants (i.e. *n* = 12). Further normality tests were used to assess the distribution of the summary group samples. For statistical comparisons between groups, paired t-tests were used in the case of normally distributed samples, and Wilcoxon signed-rank tests were used otherwise. The following effect size metrics are reported: for t-tests, the Cohen’s *d* metric; and for Wilcoxon signed-rank tests, the common language effect size (CLES). All statistical analyses were performed in Python using the Pingouin package^47^.

## Results

### Offline decoding performance assessment

Representative offline predictions of the positions of the five DOAs of the prosthetic hand are shown in Figure 2A for an able-bodied and 2B for an amputee participant, respectively. Both graphs show finger trajectories (i.e. normalised positions) in the held-out testing dataset, which comprised two repetitions of each of the nine training exercises (Supplementary Figure S1).

**Figure 2.**
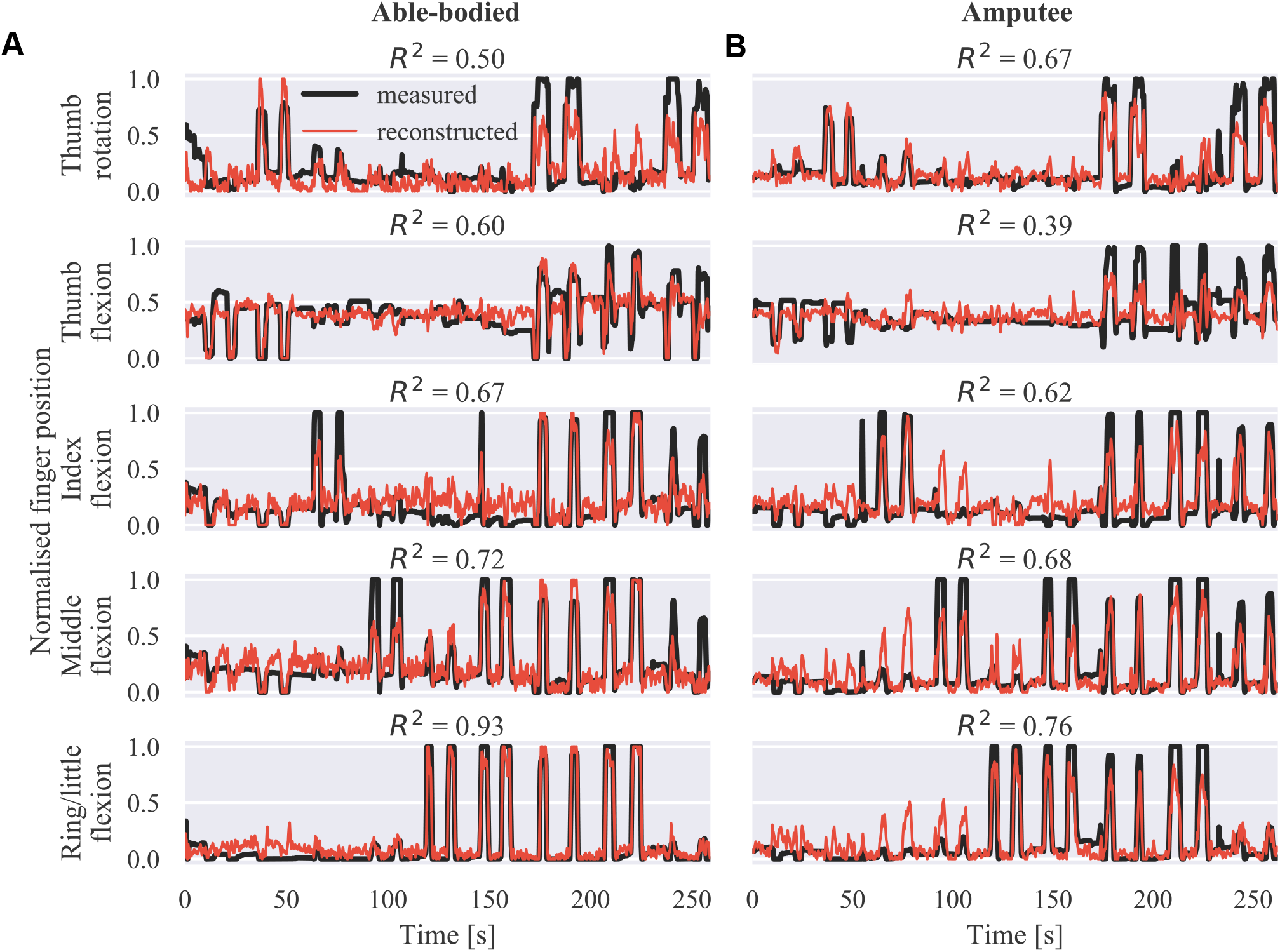
Offline finger position reconstruction. Sample test set predictions shown for the five DOAs of the robotic hand for (**A**) an able-bodied and (**B**) an amputee subject. The shown segments of activity correspond to two repetitions of each of the movements included in the training set, in the following order: thumb flexion/extension, thumb abduction/adduction, index flexion/extension, middle flexion/extension, ring/little flexion/extension, index pointer, cylindrical grip, lateral grip, and tripod grip. *R*^2^, coefficient of determination.

Offline reconstruction accuracy results are summarised in Figure 3. The multivariate R^2^ is shown in Figure 3A for all participants on the three collected datasets, that is, the training, validation, and test sets. As was to be expected, performance on the validation and test sets was slightly inferior to that on the training set. The average offline decoding accuracies in the three datasets were: training set, median 0.72, range 0.19; validation set, median 0.59, range 0.27; test set, median 0.63, range 0.22. Figure 3B shows the test set offline accuracy for individual DOAs and participants. The highest average decoding accuracy was achieved for the ring/little fingers DOA (median 0.73, range 0.41) followed by the middle finger DOA (median 0.71, range 0.24). The worst performance was observed for the thumb flexion DOA (median 0.44, range 0.45). This pattern was observed in four out of 12 participants. Finally, an overall summary is provided in Figure 3C, separately for the able-bodied and amputee groups. The average accuracy scores for the two groups were: able-bodied, median 0.63, range 0.22, *n* = 10; amputee, median 0. 60, range 0.12, *n* = 2; *n* refers to number of participants in each group.

**Figure 3.**
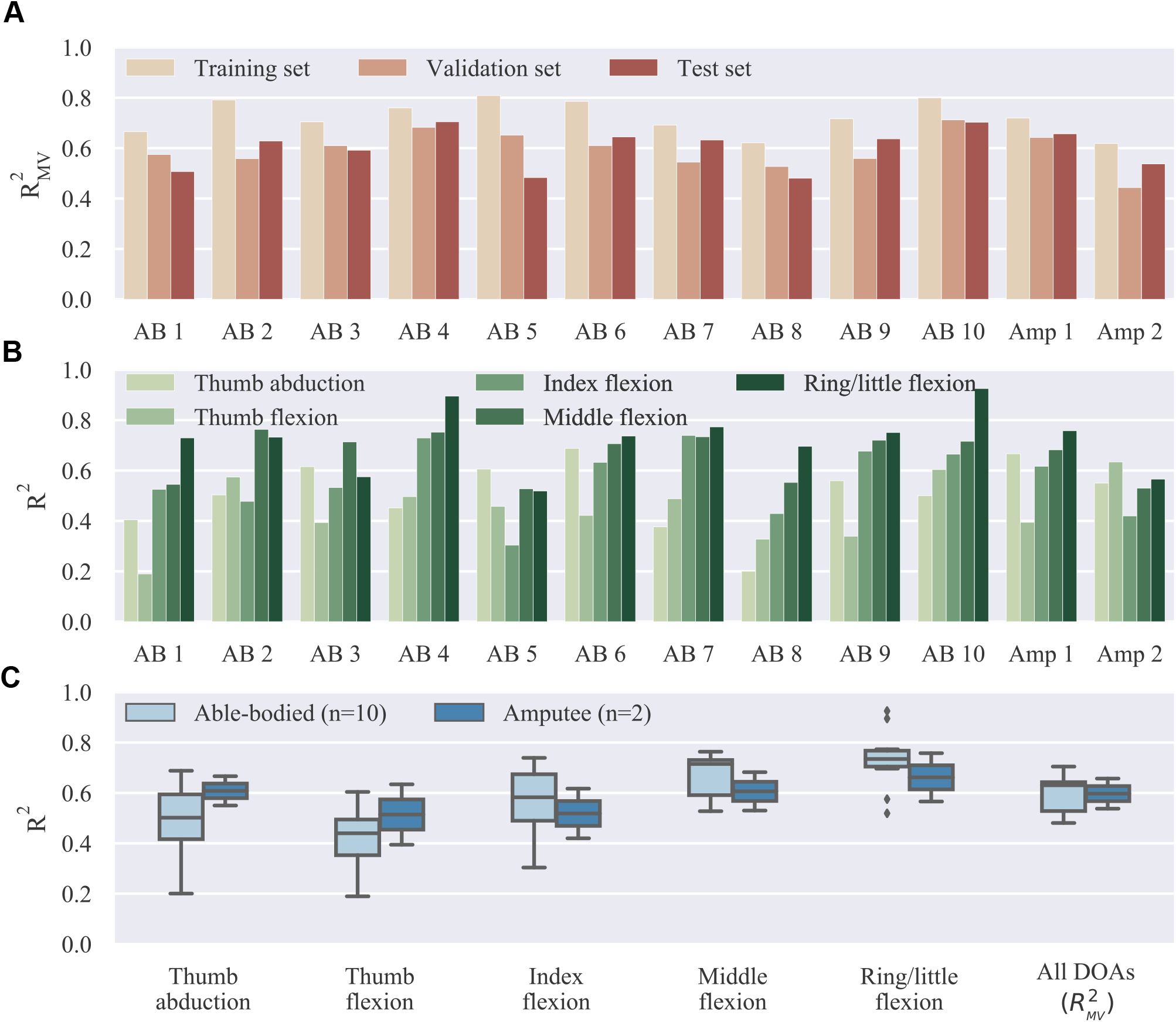
Offline decoding results summary. (**A**) Overall reconstruction accuracy on different datasets (training, validation, and test sets) for individual subjects. The three datasets were used, respectively, for model fitting, hyper-parameter optimisation, and final testing. (**B**) Offline reconstruction accuracy for individual DOAs and participants on test set. (**C**) Summary of reconstruction accuracy on test set for individual DOAs and comparison between able-bodied and amputee participants. Bars, means; straight lines, medians; solid boxes, interquartile ranges; whiskers, overall ranges of non-outlier data; diamonds, outliers; AB, able-bodied; Amp, amputee.

### Real-time experiment results

We start our analysis of the real-time control experiment by summarising the overall performance in Figure 4. We report here MAE scores between target and performed postures. A similar analysis of the closely related performance scores presented to participants at the end of the trials is provided in the Supplementary Material (Supplementary Results section). Not surprisingly, glove control performance was significantly higher than EMG control (*p* = 10^-9^, *d* = 6.5, *n* = 12; paired t-test; *n* refers to total number of participants). The median MAE scores across all participants, blocks, and trials were 0.24 (range 0.90) and 0.11 (range 0.59) for EMG and glove control, respectively. The average MAE scores across all blocks for the two groups were: EMG control, able-bodied, median 0.23, range 0.90, *n* = 1080; EMG control, amputee, median 0.26, range 0.74, *n* = 216; glove control, able-bodied, median 0.11, range 0.58, *n* = 1080; glove control, amputee, median 0.12, range 0.38, *n* = 216; *n* refers to number of single trials within each participant group and control mode.

**Figure 4.**
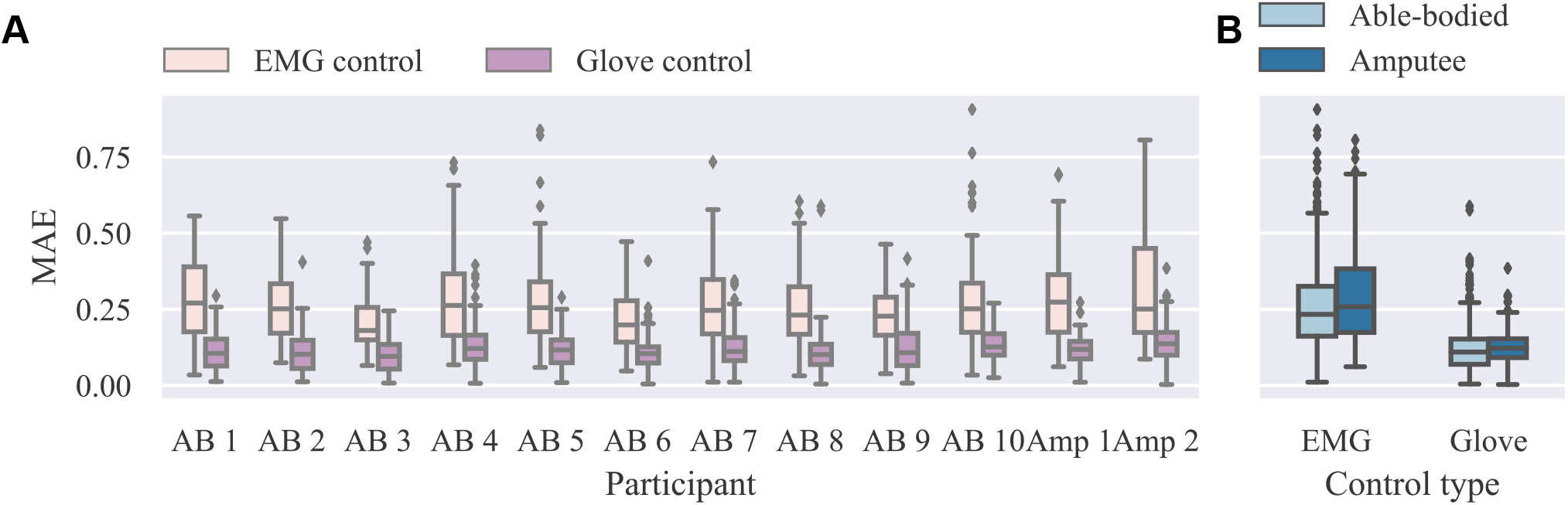
Summary results for real-time experiment. (**A**) Individual MAE scores for EMG and glove control modes. Lower scores indicate better performance. (**B**) Summary results for able-bodied and amputee groups for the two control modes. Data from all trials and postures are shown (*n* = 108 for each participant and control mode). MAE, mean absolute error; EMG, electromyography.

We now turn our attention to the effect of user practice on performance during real-time finger control. Learning curves for the real-time task are presented in Figure 5, where average performance scores are plotted against the experimental block number (ranging from one to six). In all cases, an improvement in performance can be observed as the block number increases (Figure 5A,B). A statistical comparison of early (i.e., 1-2) versus late (i.e., 5-6) blocks is provided in Figure 5C, separately for each control mode. For this analysis, able-bodied and amputee participants have been grouped together. For both control modes, average MAEs were significantly lower in late than in early blocks (EMG control, *p* = 0.02, *d* = 0.612; glove control, *p* = 10^-5^, *d* = 2.49, paired t-tests, *n* = 12; *n* hereafter refers to total number of participants). A one-to-one comparison of performance in early versus late blocks is shown in Figure 5D, where each point in the scatter plot corresponds to a single participant and control mode. For EMG control, the performance was higher in late blocks for nine out of 12 participants (one out of two amputees). For glove control, the performance in late blocks was consistently improved for all 12 participants (some points in the plot are overlaid and therefore not visible). Two videos showing one amputee participant performing the first and last blocks of the real-time posture matching task are provided as Supplementary Material (Supplementary Movies S1 and S2).

**Figure 5.**
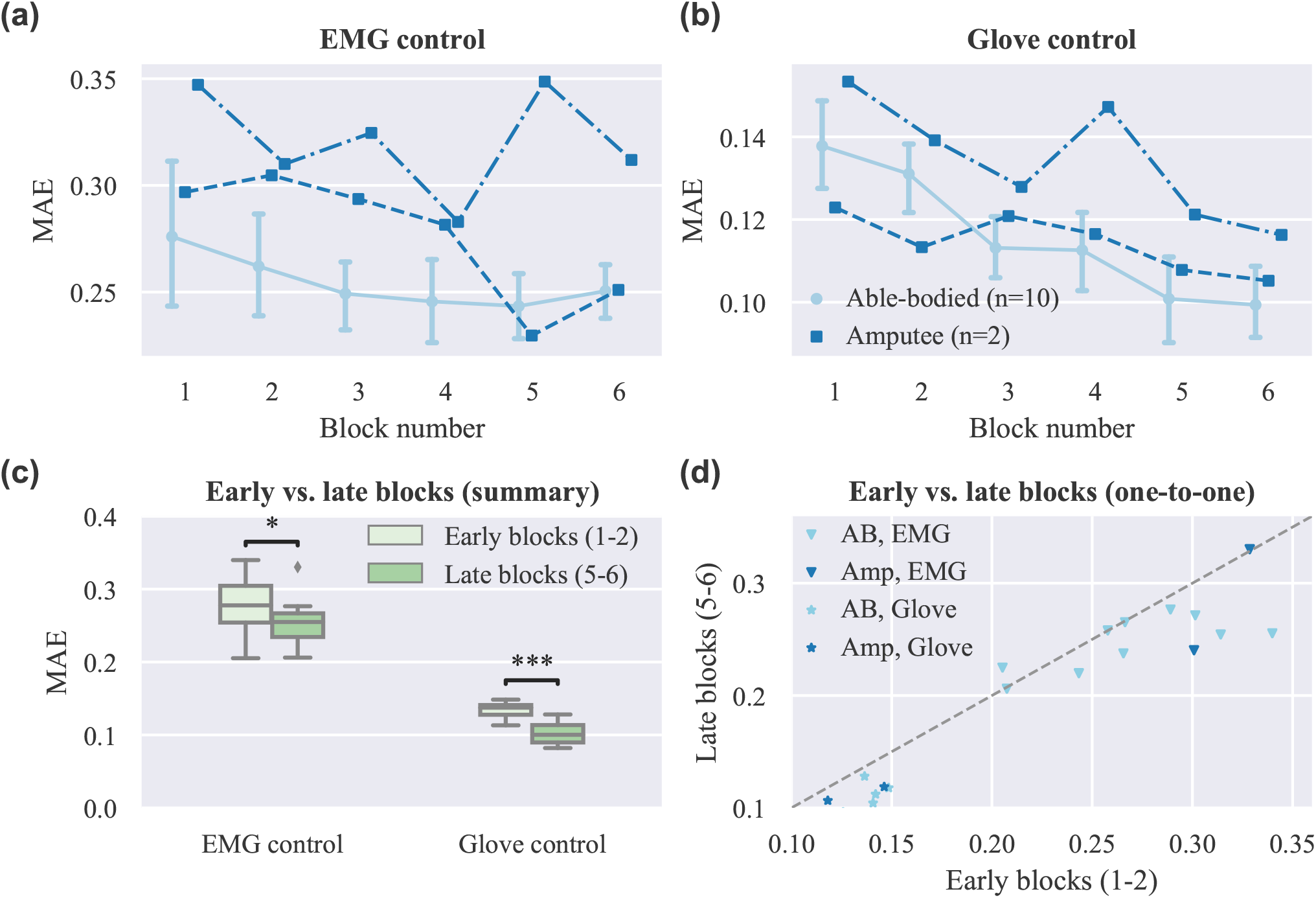
Effect of user practice on task performance (real-time experiment). (**A**)-(**B**) MAE scores are plotted against the experimental block number for (**A**) EMG and (**B**) glove control (note different y axis scales). Data for able-bodied participants are presented as means with confidence intervals. Amputee data are shown separately for each of the two individuals. (**C**) Comparison of early versus late blocks for grouped participants (able-bodied and amputee). Each block lasted 3 minutes and participants were given a 1-minute rest after block 3, therefore blocks 1 and 5 were approximately 13 minutes apart. Data shown correspond to subject averages across blocks for all participants (i.e. 10 able-bodied and two amputees). Within each block, participants replicated 18 hand postures presented to them exactly once in a pseudo-randomised order. (**D**) One-to-one comparisons of early versus late blocks averages for all participants. Each point in the scatter plot corresponds to a single participant and control mode. Points, means; error bars, 95% confidence intervals estimated via bootstrapping (1000 iterations); single asterisk, *p* < 0.05; triple asterisk, *p* < 0.001.

Next, we seek to investigate whether user practice can have an effect on the user’s muscular activity. As a first step, we perform dimensionality reduction on the recorded EMG envelopes using PCA. The top row of Figure 6 shows the cosine similarities of the first (Figure 6A) and second (Figure 6B) PCs between the first and subsequent blocks. The variance explained by the first (Figure 6C) and first two (Figure 6D) PCs extracted in block 1 is plotted against the block number in the second row of Figure 6. In the bottom row of the Figure (6E,F), the percentage of variance explained in each block is shown again against the block number, but this time the PCs used were estimated in the same blocks. For both participant groups, a decrease in similarity between the PCs in the first and subsequent blocks is observed as the block number increases. Similarly, a consistent decreasing trend is observed for the percentage of explained variance by the first two PCs estimated in the first block of trials. When using PCs extracted from the same block, the percentage of variance explained is comparable across blocks.

**Figure 6.**
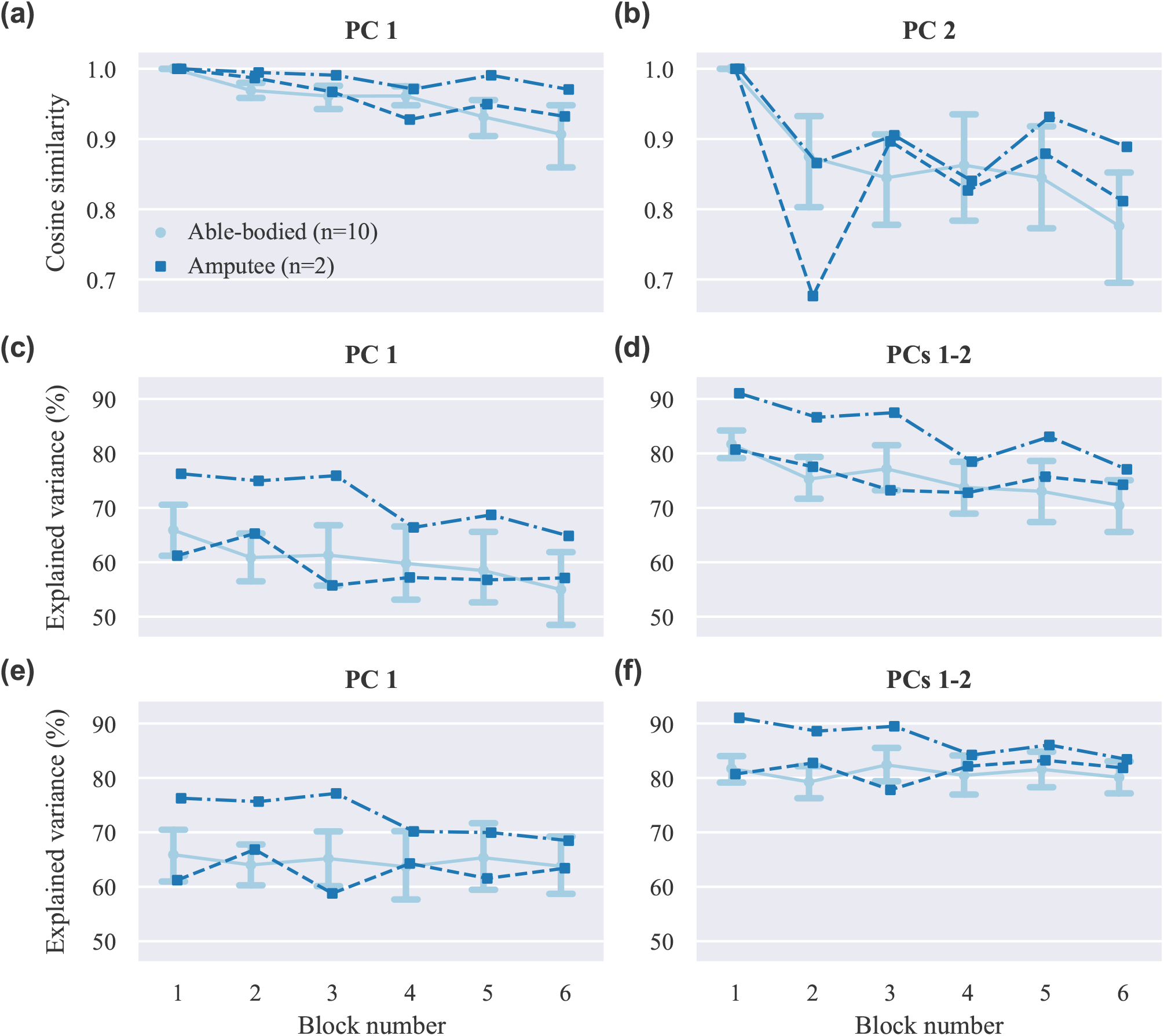
Dimensionality reduction analysis. (**A**)-(**B**) Average cosine similarities between (**A**) first (**B**) and second PCs in each block with the respective PCs computed in block 1. (**C**)-(**D**) Percentage of explained variance in each block by the (**C**) first and (**D**) first two PCs computed in block 1. (**E**)-(**F**) Percentage of explained variance in each block by (**E**) first and (**F**) first two PCs computed in the same block. PC, principal component. Data for able-bodied participants are presented as means with confidence intervals. Amputee data are shown separately for each of the two individuals.

Additionally, we compare the power of the recorded surface EMG channels across experimental blocks. This analysis is performed separately for the set of electrodes used for real-time decoding and the non-used set. The results of this analysis are presented in Figure 7A. For both groups of electrodes we observe a small, but not significant, decrease in median EMG signal power between early and late trials (used electrodes, *p* = 0.11, *CLES* = 0.632; non-used electrodes, *p* = 0.12, *CLES* = 0.636; *n* = 12 in both cases, Wilcoxon signed-rank tests). Similarly, we assess the effect of user practice on the variability of the controllable DOAs, that is, the robotic hand finger positions. Variability is assessed in terms of standard deviation during the evaluation phase of the posture matching task. The results of this analysis are presented in Figure 7C separately for EMG and glove control. For EMG control, a significant decrease in finger position variability between early and late blocks is observed (*p* = 0.01, *CLES* = 0.778). Conversely, for glove control, there is no difference between early and late blocks (*p* = 0.84, *CLES* = 0.556, *n* = 12, Wilcoxon signed-rank tests). One-to-one comparisons of average EMG power and finger position variability between early and late blocks are shown in Figure 7C and 7D, respectively, where each point in the scatter plots corresponds to a single participant and decoding condition.

**Figure 7.**
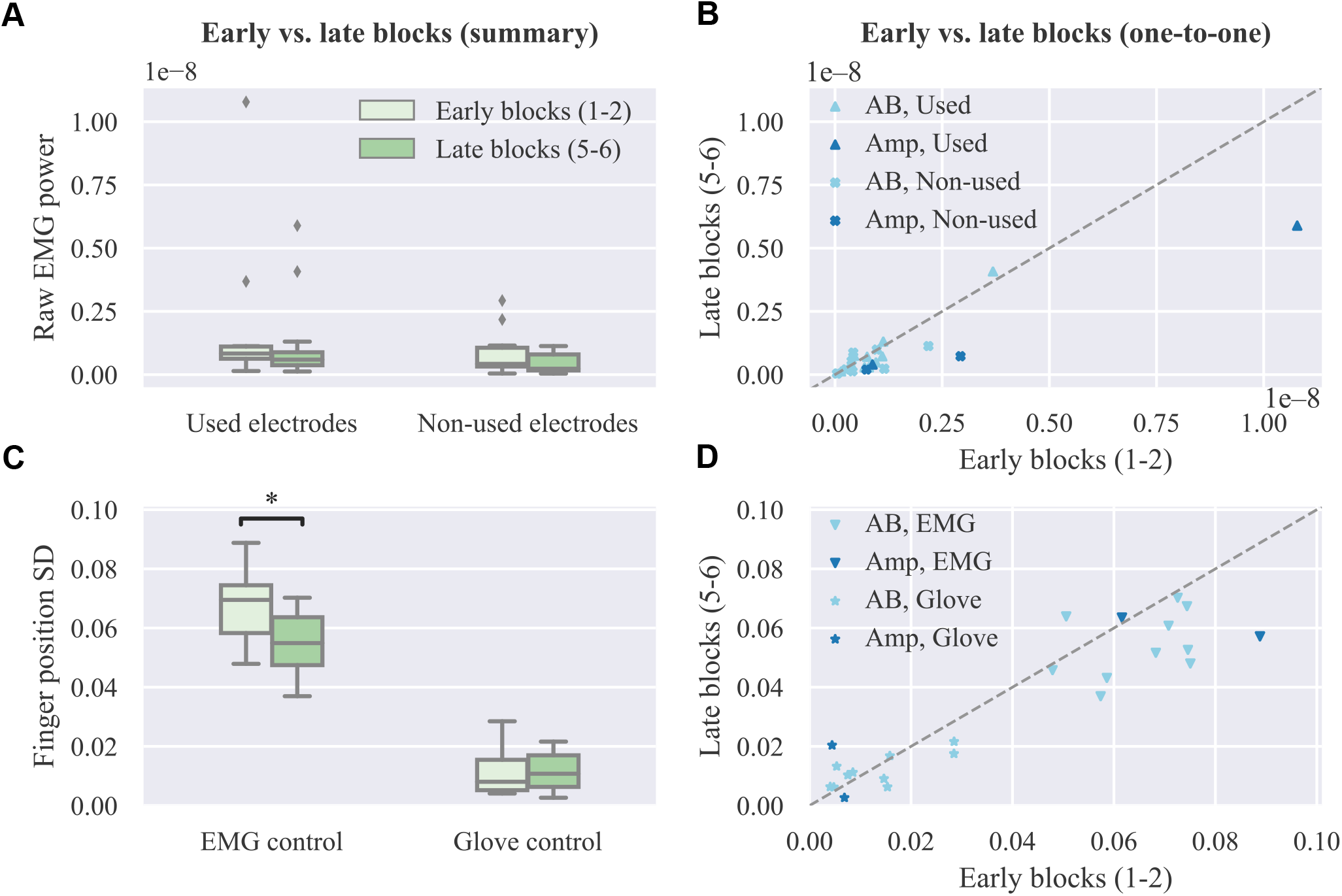
Effect of user practice on muscle force and output variability. (**A**) Average raw EMG power of used and non-used electrodes for real-time decoding in early versus late blocks. (**B**) One-to-one comparison of raw EMG power between early and late blocks. (**C**) Average finger position variability during the evaluation phase of the real-time posture matching task in early versus late blocks for EMG and glove control. (**D**) One-to-one comparison of finger position variability between early and late blocks. SD, standard deviation.

Finally, we investigate whether offline decoding accuracy can provide a reliable predictor of real-time control performance. For this reason, we compute the average MAE for each subject across all trials and blocks and compare this metric to the respective offline reconstruction accuracy score for the same subject on the test set. The results of this analysis are presented in Figure 8, where each point in the plot corresponds to a single participant. A very weak, non-significant (*p* = 0.69, *n* = 12) negative correlation is observed between offline reconstruction accuracy (i.e. multivariate R^2^) and average real-time error (i.e., MAE). Based on this observation, we conclude that it is not possible to predict the performance of real-time finger position control solely based on offline accuracy scores.

**Figure 8.**
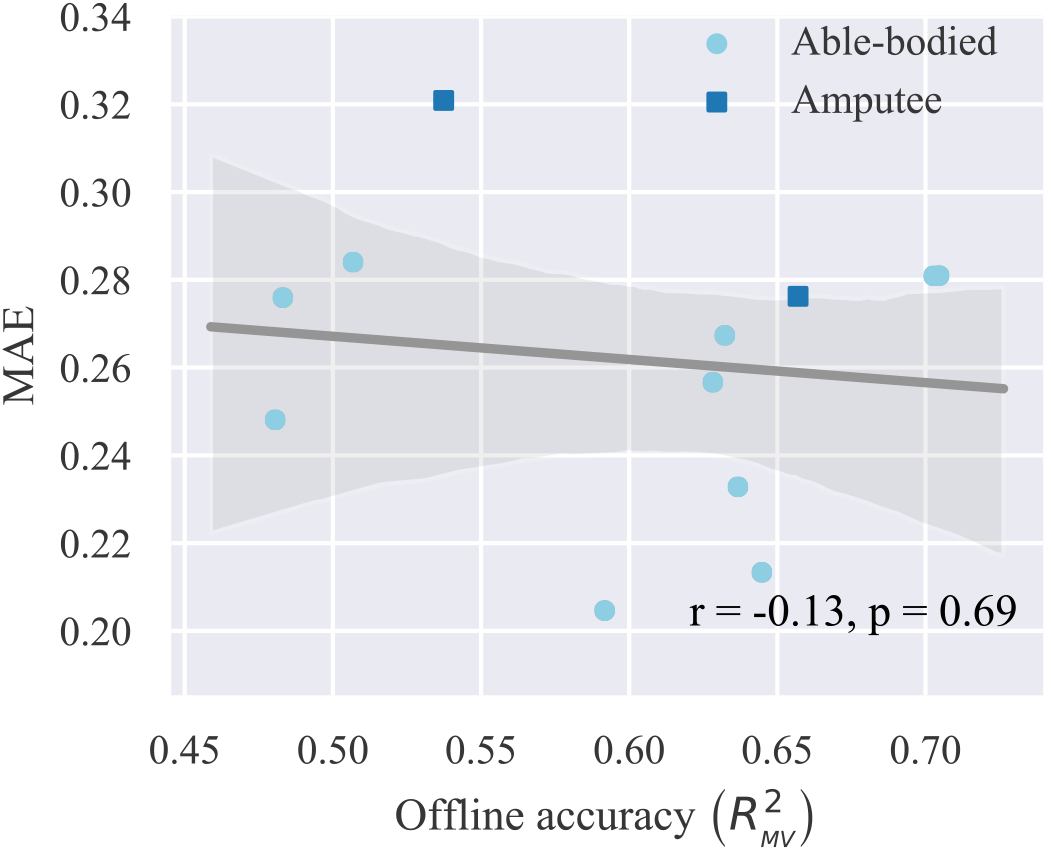
Relationship between offline reconstruction accuracy (multivariate R^2^) and real-time performance (MAE). Circles/squares, individual observations (i.e., one for each participant); line, linear regression fit; translucent band, 95% confidence intervals estimated via bootstrapping (1000 iterations); *r*, correlation coefficient; *p*, significance value.

## Discussion

The goal of this study was to investigate the effect of user practice on performance during intuitive, individual finger prosthesis control. A large body of previous work has shown that controlling a prosthesis using a non-intuitive interface, such as two-site EMG mode switching, requires motor skills that can be developed via frequent interaction with the device^2,3^. With regard to proportional, that is, continuous myoelectric control, there has been evidence that experience can lead to formation of novel, task-specific muscle synergies when the association between muscle co-activations and the DOFs of the output device is non-intuitive from a physiological perspective^31,32,36,48^. Therefore, non-intuitive paradigms may require training before a user is able to control a prosthesis at its full capacity. On the other hand, the use of more intuitive interfaces, such as those based on multi-site EMG signal classification, can alleviate some of this burden due to relying on a natural association between muscle contractions and prosthesis activations. Previous work has shown that even in the case of intuitive interfaces, user practice results in substantial control performance improvement^5–8^. However, these studies were concerned with classification-based control, which still lacks complete intuitiveness due to the discrete and sequential nature of the hand actuation mechanism. Here, we investigated the effect of user adaptation on myoelectric control when using a decoder mapping EMG features onto prosthetic digit positions.

We found evidence that the performance of intuitive, independent prosthesis finger control can benefit from user experience gathered during real-time, closed-loop interaction with the control interface. In our experiment, two types of feedback were provided, namely, visual, since the prosthetic hand was within the visual field of the participant and responded to their control input, and a performance score, which was presented to the participants at the end of each trial. We hypothesised that despite the intuitiveness of the controller, experience should allow users to improve their performance. Indeed, we observed a significant decrease in target posture mismatch within approximately 20 minutes of interaction with the prosthesis (Figure 5).

Of particular interest is the question of whether the observed improvement in performance can be retained during long-term use. Previous work has demonstrated an increase in classification-based myoelectric performance after a 6-8 week home trial^8^. Whether a similar pattern can be observed with individual finger control remains to be investigated. Furthermore, it shall be compelling to investigate whether long-term performance improvement is accompanied by permanent changes in forward neuromotor control (i.e., motor learning). Our study has demonstrated that users can adapt rather quickly to improve performance on a specific task based on feedback. However, to assess long-term adaptation, a more extended study spanning across multiple sessions and testing generalisation on novel tasks might be needed^49^.

The dimensionality reduction analysis (Figure 6A,B) revealed that over the course of our experiments, substantial changes occurred in the covariance structure of the recorded EMG signal envelopes, and therefore the direction and variance explained by the first two PCs (Figure 6C,D). Although such temporal changes in muscle co-activation patterns might in part reflect the non-stationary nature of surface EMG recordings, when combined with the observed increase in task performance, these changes may be primarily mediated by user adaptation in muscle recruitment. It is worth mentioning that although the extracted PCs reflect muscle co-activation patterns, they do not directly correspond to muscle synergies, as we did not target specific muscles during electrode positioning. However, by taking into consideration that the EMG electrodes record in this setting a superposition of the activity of different muscles, and also given that PCA produces a linear transformation of the input space, it is reasonable to expect that similar results would have been obtained had we targeted specific muscles. It is also worth noting that the dimensionality reduction analysis was performed on the EMG envelopes, that is, the mean absolute value of the recorded signals, whereas various non-linear feature transformations were used for decoding finger positions from EMG signals. Therefore, there is no direct correspondence between the estimated PCs and the principal directions of the regression problem^44^, which has a substantially higher dimensionality. It can be observed in Figure 6C,D that the first two PCs explained a higher percentage of the overall variance in the two amputee than in the able-bodied participants. We attribute this to the smaller number of electrodes used in the former case (i.e. 12 and 13 electrodes for amputees as opposed to 16 electrodes for able-bodied participants). The percentage of explained variance in each block by the PCs computed in the same block remained constant during the experiment (Figure 6E,F). This observation rules out the possibility that the decrease in explained variance by the PCs extracted in the first block (Figure 6C,D) is due to exogenous parameters, hence further suggesting that this reduction is caused by changes in muscle co-activation patterns emerging from short-term user adaptation.

Force field adaptation studies have previously shown that humans learn to optimise limb impedance to minimise metabolic costs and movement error simultaneously^50^. In addition to a decrease in movement error, we also observed a small, however non-significant, reduction in overall EMG power exerted by the participants’ muscles and a significant decrease in the variability of the controllable DOAs (Figure 7). Both of these observations are compatible with the notion of limb impedance optimisation. Notably, we did not observe a decrease in finger position variability with glove control, hence implying that the respective reduction with EMG control should be indeed attributed to changes in the recorded muscle signals. A key difference between our study and previous work50 is that the user and the device were not mechanically linked, but in both cases the effector was unstable and with practice subjects learned to enhance its stability. The decrease in median EMG power was non-significant (*p* = 0.10) and therefore no definitive conclusions can be drawn regarding whether and how user training can lead to a reduction in muscle metabolic cost during myoelectric control. Future experiments with amputee participants wearing a prosthetic hand can reveal the extent to which energy-efficient control can be achieved. It is unlikely that the observed trend is due to muscle fatigue, since it is known that the latter is associated with an increase in EMG power with a simultaneous reduction of median frequency of the EMG spectrum^51,52^.

A previous study that made use of a performance score that was similar to the one presented to the participants at the end of the trials reported an increase from 0% to 40% after approximately 200 trials32. In the current study, the average performance with EMG control increased from 33.48% to 36.50% corresponding to a decrease in average MAE from 0.28 to 0.26 (Figure 5A) after 108 trials. Although this improvement is smaller than the one reported previously, this finding should not be surprising; the previous study used a pre-determined, fixed, and non-intuitive mapping from muscle activity to the DOAs of the prosthetic hand, and thus, participants had to learn the underlying control principle, that is, an inverse model of the interface30 from scratch. Conversely, in our study, the mappings were based on regression models trained with user-specific data; hence, the mappings were intuitive for all participants and the baseline performance at the start of the experiment was well above zero.

Our results agree with previous work suggesting that experience can help humans improve their performance at myoelectric control tasks, even in the case of intuitive interfaces^48^. A question that may naturally arise is why one should expect such improvement when using biomimetic, intuitive myoelectric decoders. Before addressing this sensible question, one should first note that in our experiments an increase in performance was also observed in the case of robotic hand teleoperation using the data glove, despite the very high level of intuitiveness of this particular task. With this important information in mind, we seek to provide the following justification: despite being intuitive from a physiological perspective, the myoelectric controller is still far from natural; this is due to many differences between the human and robotic hands, including, but not limited to, the number of DOFs, the finger anatomical structure, and the range of finger movement. Furthermore, our control algorithm mapped recorded muscle activity onto finger joint positions without taking into account joint velocities or digit forces which, in the human body, are also controlled by muscle contractions. In the case of EMG control, although decoding accuracy was relatively high, it was still far from perfect (Figure 3). Therefore, it is likely that participants performed compensatory contractions to correct for prediction errors^12,40^. Finally, it is worth noting that the lack of proprioception in amputees can only exacerbate the challenges outlined above. Taking everything into consideration, it is clear that despite our best efforts, it is still impossible to develop biomimetic myoelectric control schemes that feel entirely natural to the user, unless all the following conditions are simultaneously met: the end device is a perfect replication of the human hand, sensing technologies and decoding algorithms allow for near-perfect reconstruction of movement intent and, finally, artificial proprioceptive information is fed back to the user. In any other case, a certain amount of adaptation is still very likely to take place during interaction with the device, which has the potential to substantially improve control performance.

In addition to an increase in control accuracy, user training may have additionally resulted in a reduction in reaction time. However, given the long preparation window, in combination with the fact that performance was only assessed with respect to the final posture and not the followed trajectories, it seems unlikely that this factor could explain the observed increase in control performance. One such example is given in Figure S5, where two trials are compared for the same participant and posture, one at the start and one at the end of the experimental session. For the shown example, after training, the participant was able to reach the desired posture in less time and with better accuracy. However, in both cases, the final posture was achieved before the start of the evaluation phase. Hence, the difference in performance scores is only due to the higher accuracy of the end posture in the late trial. The length of the preparation window was set during pilot trials to the chosen value (i.e., 3.5 s), as this was found to offer a good trade-off between the desired task difficulty and the ability to assess performance without being influenced by a potential decrease in user reaction time.

In the context of myoelectric classification, it has been previously shown that a discrepancy exists between offline accuracy and real-time control performance^38^. With regard to continuous wrist control, it has been shown that only a weak correlation exists between offline R^2^ and metrics characterising real-time performance during a target achievement control test, such as completion rate, completion times, overshoots, throughput, speed, and efficiency coefficient^12^. To assess whether a similar statement could be made about continuous finger position control, we compared offline reconstruction accuracy to performance scores during the real-time posture matching task. In agreement with previous work^12,38^, a very weak, non-significant correlation was found between offline accuracy and real-time performance. Such differences between offline decoding and real-time control, which may be primarily attributed to user adaptation taking place during close-loop interaction, further reiterate the need for testing prosthetic control methodologies with real-time experiments^12,38,39,53^.

In this study, we focused on non-invasive, continuous position control of individual digits. In line with previous work^20^, we have shown that it is feasible, in principle, to use surface EMG measurements from the forearm of able-bodied and transradial amputee subjects to decode finger positions and subsequently use these estimates to control the individual digits of a prosthesis in real-time. The set of controllable DOAs included flexion of all fingers and thumb opposition. The ring and little fingers were controlled together because of mechanical coupling in the robotic device used in our experiments. Offline analysis revealed that thumb movement (flexion and rotation) was the most challenging to decode (Figure 3). This is not surprising, given that thumb muscles are either intrinsic or located in the distal part of the forearm. In this work, however, we focused on transradial amputation and, therefore, recorded EMG activity from the proximal part of the forearm only.

The continuous finger position controller has two main advantages: intuitiveness and dexterity. As has already been pointed out, training regression models using muscle signals and glove data recorded from the end-user creates an intuitive association between muscle activity and finger movement and, thus, the user does not need to learn a new mapping from scratch. Dexterity naturally arises from the fact that the user can control individual digits in a continuous space. One particular advantage of this scheme over discrete control schemes, for example, classification-based grip selection, is the ability to move from one type of grip to another without the need for executing an intermediate hand opening action. The high level of dexterity, however, comes at a price; decoding independent finger movement is a much more challenging task than classifying EMG activity into hand postures. In its current form, the proposed scheme is unlikely to be suitable for clinical adoption by amputees, as significant improvements are required to ensure its long-term viability. For example, one simplification made in this study was that the posture of the participants’ forearm was kept fixed throughout the experimental sessions. This simplification would not occur in a realistic scenario and, thus, it is expected that performance would deteriorate due to the limb position effect^54^. Nevertheless, given the potential of this method to achieve intuitive and truly dexterous prosthetic control, we consider it is worthwhile pursuing further research in this direction.

It is worth mentioning that an invasive approach might indeed be required to achieve robust finger position control. Intramuscular EMG recordings have been previously used for continuous finger control of both virtual23 and robotic24 hands. Both of these studies, however, had the following two limitations: firstly, they used a one-to-one mapping from individual pairs of muscles to DOAs of the hand; secondly, they were limited to able-bodied participants. An alternative avenue would be to investigate the use of multivariate regression models in mapping the activity of multiple muscles onto prosthesis DOAs, as opposed to the previously used one-to-one mapping schemes. Another compelling possibility would be to test the performance of continuous finger control in patients having undergone targeted muscle reinnervation. It has been previously demonstrated that hand/wrist movements can be classified with high accuracy in targeted muscle reinnervation patients^55^. Whether robust individual finger control can be achieved using a similar invasive approach remains to be investigated.

As a final note, we seek to re-emphasise the important role that user adaptation could play in myoelectric control of prosthetic fingers, regardless of the origin of control signals. We have shown here that even with an intuitive decoder, humans can improve their performance in a biofeedback myoelectric task within a short period of time. In line with previous reports from the myoelectric classification and wrist control literature^12,38^, we conclude that future efforts should focus on putting the human in the loop and evaluating control methodologies with real-time, closed-loop experiments. We firmly believe that further advancements can be achieved by explicitly taking into account the remarkable plasticity of the human brain when designing myoelectric control interfaces.

## Conflict of Interest Statement

The authors declare that the research was conducted in the absence of any commercial or financial relationships that could be construed as a potential conflict of interest.

## Author Contributions

AK designed and implemented the experimental protocol. AK and KN conducted the experiments. AK analyzed the data. All authors contributed to manuscript preparation. All authors read and approved the final manuscript.

## Funding

AK was supported in part by grants EP/F500386/1 and BB/F529254/1 for the University of Edinburgh School of Informatics Doctoral Training Centre in Neuroinformatics and Computational Neuroscience from the UK Engineering and Physical Sciences Research Council (EPSRC), UK Biotechnology and Biological Sciences Research Council (BBSRC), and the UK Medical Research Council (MRC). AK is currently supported by EPSRC grant EP/R004242/1. SV is supported by the Microsoft Research RAEng. Fellowship and EPSRC grant EP/L016834/1. KN is supported by EPSRC grants EP/M025977/1, EP/N023080/1, and EP/R004242/1.

## Acknowledgments

The authors are grateful to the two amputee volunteers for taking part in the study. The authors would like to thank Matthew Dyson for useful discussion.

## Footnotes

1 https://www.coaptengineering.com/

## References

1. Farina, D. et al. The extraction of neural information from the surface EMG for the control of upper-limb prostheses: emerging avenues and challenges. IEEE Trans. Neural Syst. Rehabil. Eng. 22, 797–809 (2014).

2. Bouwsema, H., van der Sluis, C. K. & Bongers, R. M. Learning to control opening and closing a myoelectric hand. Arch. Phys. Med. Rehabil. 91, 1442–1446 (2010).

3. Clingman, R. & Pidcoe, P. A novel myoelectric training device for upper limb prostheses. IEEE Trans. Neural Syst. Rehabil. Eng. 22, 879–885 (2014).

4. Herberts, P., Almström, C., Kadefors, R. & Lawrence, P. D. Hand prosthesis control via myoelectric patterns. Acta Orthop. Scand. 44, 389–409 (1973).

5. Bunderson, N. E. & Kuiken, T. A. Quantification of feature space changes with experience during electromyogram pattern recognition control. IEEE Trans. Neural Syst. Rehabil. Eng. 20, 239–246 (2012).

6. Powell, M. A., Kaliki, R. R. & Thakor, N. V. User training for pattern recognition-based myoelectric prostheses: improving phantom limb movement consistency and distinguishability. IEEE Trans. Neural Syst. Rehabil. Eng. 22, 522–532 (2014).

7. He, J. et al. User adaptation in long-term, open-loop myoelectric training: implications for EMG pattern recognition in prosthesis control. J. Neural Eng. 12, 046005 (2015).

8. Hargrove, L. J., Miller, L. A., Turner, K. & Kuiken, T. A. Myoelectric pattern recognition outperforms direct control for transhumeral amputees with targeted muscle reinnervation: a randomized clinical trial. Sci. Rep. 7, 13840 (2017).

9. Fougner, A., Stavdahl, Ø., Kyberd, P. J., Losier, Y. G. & Parker, P. A. Control of upper limb prostheses: terminology and proportional myoelectric control–A review. IEEE Trans. Neural Syst. Rehabil. Eng. 20, 663–677 (2012).

10. Muceli, S., Jiang, N. & Farina, D. Extracting signals robust to electrode number and shift for online simultaneous and proportional myoelectric control by factorization algorithms. IEEE Trans. Neural Syst. Rehabil. Eng. 22, 623–633 (2014).

11. Hahne, J. M. et al. Linear and nonlinear regression techniques for simultaneous and proportional myoelectric control. IEEE Trans. Neural Syst. Rehabil. Eng. 22, 269–279 (2014).

12. Jiang, N., Vujaklija, I., Rehbaum, H., Graimann, B. & Farina, D. Is accurate mapping of EMG signals on kinematics needed for precise online myoelectric control? IEEE Trans. Neural Syst. Rehabil. Eng. 22, 549–558 (2014).

13. Smith, L. H., Kuiken, T. A. & Hargrove, L. J. Real-time simultaneous and proportional myoelectric control using intramuscular EMG. J. Neural Eng. 11, 066013 (2014).

14. Afshar, P. & Matsuoka, Y. Neural-based control of a robotic hand: evidence for distinct muscle strategies. In Proc. IEEE Int. Conf. Robot. Autom. (ICRA), vol. 5, 4633–4638 (2004).

15. Smith, R. J., Tenore, F., Huberdeau, D., Etienne-Cummings, R. & Thakor, N. V. Continuous decoding of finger position from surface EMG signals for the control of powered prostheses. In Proc. IEEE Int. Conf. Eng. Med. Biol. Soc. (EMBC), 197–200 (2008).

16. Ngeo, J. G., Tamei, T. & Shibata, T. Continuous and simultaneous estimation of finger kinematics using inputs from an EMG-to-muscle activation model. J. NeuroEng. Rehabil. 11, 122 (2014).

17. Krasoulis, A., Vijayakumar, S. & Nazarpour, K. Evaluation of regression methods for the continuous decoding of finger movement from surface EMG and accelerometry. In Proc. IEEE/EMBS Int. Conf. Neural Eng. (NER), 631–634 (2015).

18. Xiloyannis, M., Gavriel, C., Thomik, A. A. C. & Faisal, A. A. Gaussian Process autoregression for simultaneous proportional multi-modal prosthetic control with natural hand kinematics. IEEE Trans. Neural Syst. Rehabil. Eng. 25, 1785–1801 (2017).

19. Smith, R., Huberdeau, D., Tenore, F. & Thakor, N. Real-time myoelectric decoding of individual finger movements for a virtual target task. In Proc. IEEE Int. Conf. Eng. Med. Biol. Soc. (EMBC), 2376–2379 (2009).

20. Cipriani, C. et al. Online myoelectric control of a dexterous hand prosthesis by transradial amputees. IEEE Trans. Neural Syst. Rehabil. Eng. 19, 260–270 (2011).

21. Ngeo, J. G. et al. Control of an optimal finger exoskeleton based on continuous joint angle estimation from EMG signals. In Proc. IEEE Int. Conf. Eng. Med. Biol. Soc. (EMBC), 338–341 (2013).

22. Birdwell, J. A., Hargrove, L. J., Kuiken, T. A. & ff Weir, R. F. Activation of individual extrinsic thumb muscles and compartments of extrinsic finger muscles. J. Neurophysiol. 110, 1385–1392 (2013).

23. Birdwell, J. A., Hargrove, L. J., ff Weir, R. F. & Kuiken, T. A. Extrinsic finger and thumb muscles command a virtual hand to allow individual finger and grasp control. IEEE Trans. Biomed. Eng. 62, 218–226 (2015).

24. Cipriani, C., Segil, J. L., Birdwell, J. A. & ff Weir, R. F. Dexterous control of a prosthetic hand using fine-wire intramuscular electrodes in targeted extrinsic muscles. IEEE Trans. Neural Syst. Rehabil. Eng. 22, 828–836 (2014).

25. Castellini, C., Gruppioni, E., Davalli, A. & Sandini, G. Fine detection of grasp force and posture by amputees via surface electromyography. J. Physiol. 103, 255–262 (2009).

26. Nielsen, J. L. G. et al. Simultaneous and proportional force estimation for multifunction myoelectric prostheses using mirrored bilateral training. IEEE Trans. Biomed. Eng. 58, 681–688 (2011).

27. Gijsberts, A. et al. Stable myoelectric control of a hand prosthesis using non-linear incremental learning. Front. Neurorobot. 8 (2014).

28. Patel, G., Nowak, M. & Castellini, C. Exploiting knowledge composition to improve real-life hand prosthetic control. IEEE Trans. Neural Syst. Rehabil. Eng. 25, 967–975 (2017).

29. Gailey, A., Artemiadis, P. & Santello, M. Proof of concept of an online EMG-based decoding of hand postures and individual digit forces for prosthetic hand control. Front. Neurol. 8 (2017).

30. Dyson, M., Barnes, J. & Nazarpour, K. Myoelectric control with abstract decoders. J. Neural Eng. 15, 056003 (2018).

31. Nazarpour, K., Barnard, A. & Jackson, A. Flexible cortical control of task-specific muscle synergies. J. Neurosci. 32, 12349–12360 (2012).

32. Pistohl, T., Cipriani, C., Jackson, A. & Nazarpour, K. Abstract and proportional myoelectric control for multi-fingered hand prostheses. Ann. Biomed. Eng. 41, 2687–2698 (2013).

33. Antuvan, C. W., Ison, M. & Artemiadis, P. Embedded human control of robots using myoelectric interfaces. IEEE Trans. Neural Syst. Rehabil. Eng. 22, 820–7 (2014).

34. Pistohl, T., Joshi, D., Ganesh, G., Jackson, A. & Nazarpour, K. Artificial proprioceptive feedback for myoelectric control. IEEE Trans. Neural Syst. Rehabil. Eng. 23, 498–507 (2015).

35. Dyson, M., Barnes, J. & Nazarpour, K. Abstract myoelectric control with EMG drive estimated using linear, kurtosis and Bayesian filtering. In Proc. IEEE/EMBS Int. Conf. Neural Eng. (NER), 54–57 (2017).

36. Ison, M. & Artemiadis, P. Proportional myoelectric control of robots: muscle synergy development drives performance enhancement, retainment, and generalization. IEEE Trans. Robot. 31, 259–268 (2015).

37. Ison, M., Vujaklija, I., Whitsell, B., Farina, D. & Artemiadis, P. High-density electromyography and motor skill learning for robust long-term control of a 7-DoF robot arm. IEEE Trans. Neural Syst. Rehabil. Eng. 24, 424–433 (2016).

38. Ortiz-Catalan, M., Rouhani, F., Brånemark, R. & Håkansson, B. Offline accuracy: a potentially misleading metric in myoelectric pattern recognition for prosthetic control. In Proc. IEEE Int. Conf. Eng. Med. Biol. Soc. (EMBC), 1140–1143 (2015).

39. Vujaklija, I. et al. Translating research on myoelectric control into clinics—Are the performance assessment methods adequate? Front. Neurorobot. 11 (2017).

40. Hahne, J. M., Markovic, M. & Farina, D. User adaptation in myoelectric man-machine interfaces. Sci. Rep. 7, 4437 (2017).

41. Krasoulis, A., Kyranou, I., Erden, M. S., Nazarpour, K. & Vijayakumar, S. Improved prosthetic hand control with concurrent use of myoelectric and inertial measurements. J. NeuroEng. Rehabil. 14, 71 (2017).

42. Boostani, R. & Moradi, M. H. Evaluation of the forearm EMG signal features for the control of a prosthetic hand. Physiol. Meas. 24, 309–319 (2003).

43. Perreault, E. J., Kirsch, R. F. & Acosta, A. M. Multiple-input, multiple-output system identification for characterization of limb stiffness dynamics. Biol. Cybern. 80, 327–337 (1999).

44. Krasoulis, A., Nazarpour, K. & Vijayakumar, S. Towards low-dimensional proportional myoelectric control. In Proc. IEEE Int. Conf. Eng. Med. Biol. Soc. (EMBC), 7155–7158 (2015).

45. Pudil, P., Novovičová, J. & Kittler, J. Floating search methods in feature selection. Pattern recognition letters 15, 1119–1125 (1994).

46. Pedregosa, F. et al. Scikit-learn: Machine learning in Python. J. Mach. Learn. Res. 12, 2825–2830 (2011).

47. Vallat, R. Pingouin: statistics in python. J. Open Source Softw. 3, 331 (2018).

48. Radhakrishnan, S. M., Baker, S. N. & Jackson, A. Learning a novel myoelectric-controlled interface task. J. Neurophysiol. 100, 2397–2408 (2008).

49. Kantak, S. S. & Winstein, C. J. Learning–performance distinction and memory processes for motor skills: A focused review and perspective. Behav. brain research 228, 219–231 (2012).

50. Burdet, E., Osu, R., Franklin, D. W., Milner, T. E. & Kawato, M. The central nervous system stabilizes unstable dynamics by learning optimal impedance. Nature 414, 446 (2001).

51. Luttmann, A., Jäger, M. & Laurig, W. Electromyographical indication of muscular fatigue in occupational field studies. Int. J. Ind. Ergonomics 25, 645–660 (2000).

52. Bartuzi, P. & Roman-Liu, D. Assessment of muscle load and fatigue with the usage of frequency and time-frequency analysis of the emg signal. Acta bioengineering biomechanics 16, 31–39 (2014).

53. Jiang, N., Dosen, S., Müller, K.-R. & Farina, D. Myoelectric control of artificial limbs –is there a need to change focus? [In the spotlight]. IEEE Signal Proc. Mag. 29, 152–150 (2012).

54. Fougner, A., Scheme, E. J., Chan, A. D. C., Englehart, K. B. & Stavdahl, Ø. Resolving the limb position effect in myoelectric pattern recognition. IEEE Trans. Neural Syst. Rehabil. Eng. 19, 644–651 (2011).

55. Kuiken, T. A. et al. Targeted muscle reinnervation for real-time myoelectric control of multifunction artificial arms. JAMA 301, 619–628 (2009).

