## Supplementary Material for "Effect of user adaptation on prosthetic finger control with an intuitive myoelectric decoder"

#### Supplementary Figures

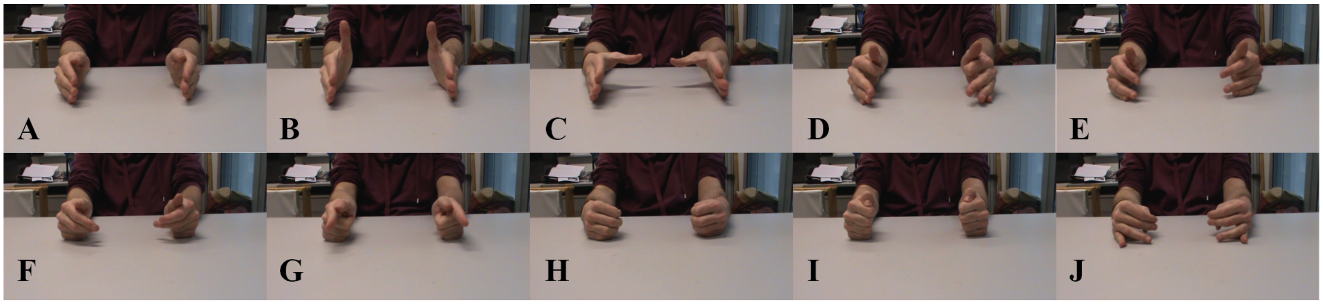


**Figure 1.** Training exercises. Subjects were instructed to perform the following bilateral mirrored movements: **(A)** rest; **(B)** thumb flexion/extension; **(C)** thumb abduction/adduction; **(D)** index flexion/extension; **(E)** middle flexion/extension; **(F)** ring/little flexion/extension; **(G)** index pointer; **(H)** cylindrical grip; **(I)** lateral grip; **(J)** tripod grip.

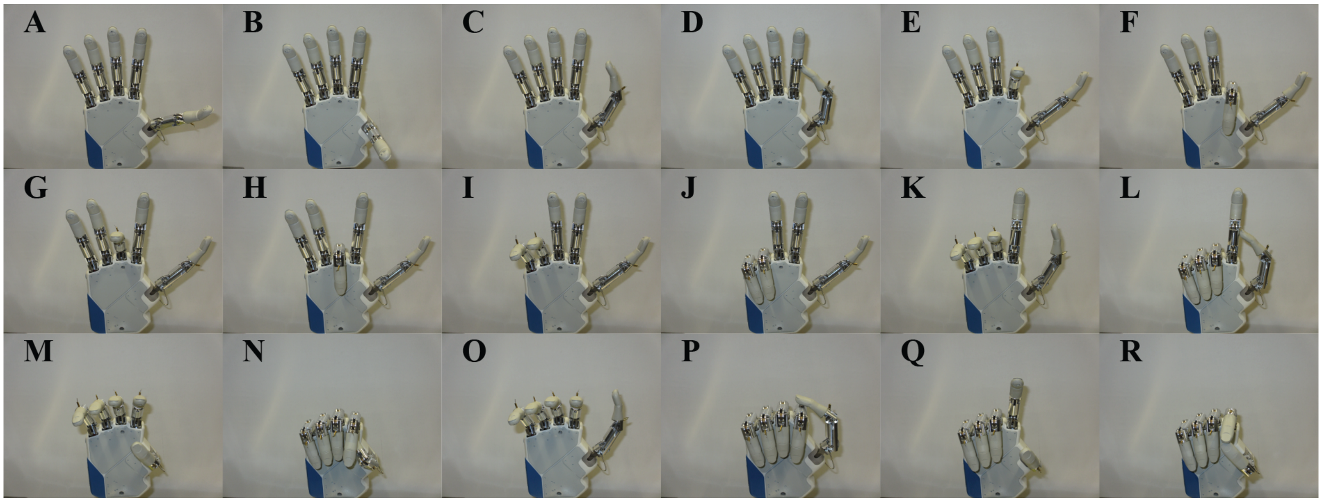


**Figure 2.** Target poses for real-time posture matching task. **(A)** Thumb abduction (half); **(B)** thumb abduction (full); **(C)** thumb flexion (half); **(D)** thumb flexion (full); **(E)** index flexion (half); **(F)** index flexion (full); **(G)** middle flexion (half); **(H)** middle flexion (full); **(I)** ring/little flexion (half); **(J)** ring/little flexion (full); **(K)** index pointer (half); **(L)** index pointer (full); **(M)** cylindrical grip (half); **(N)** cylindrical grip (full); **(O)** lateral grip (half); **(P)** lateral grip (full); **(Q)** pinch grip (half); **(R)** pinch grip (full).

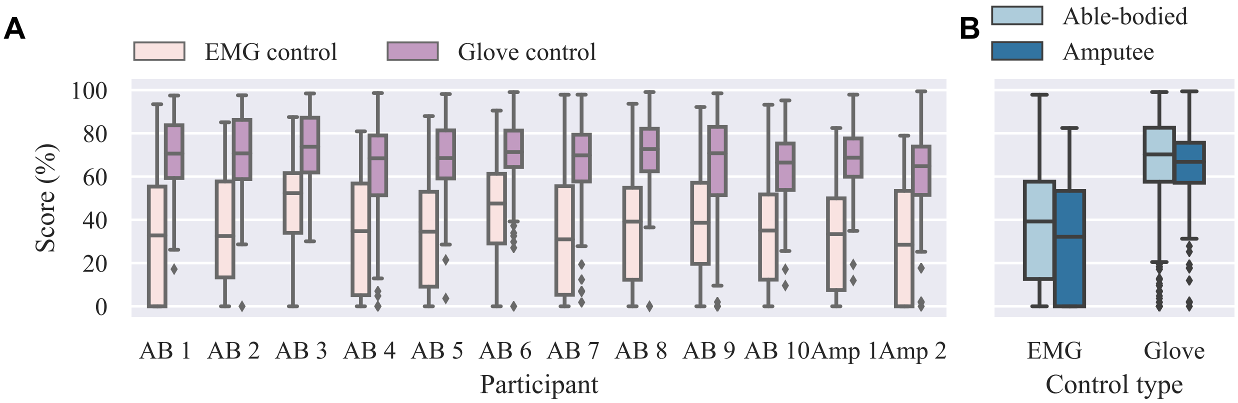


**Figure 3.** Summary results for real-time experiment (performance scores). **(A)** Performance scores received by participants after trial end shown for individual subjects, separately for EMG and glove control modes; **(B)** summary scores for able-bodied and amputee groups for the two control modes. Straight lines, medians; solid boxes, interquartile ranges; whiskers, overall ranges of non-outlier data; diamonds, outliers; AB, able-bodied; Amp, amputee.

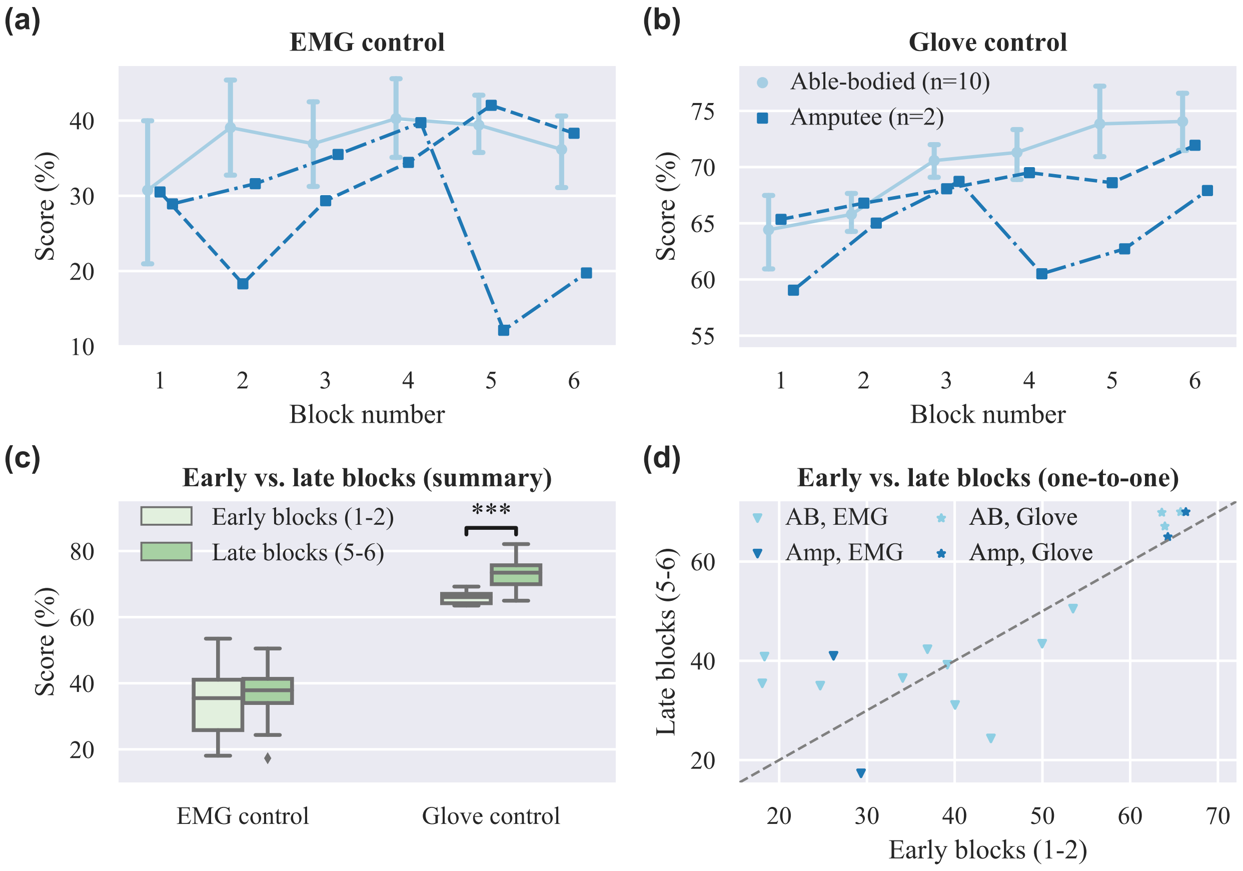


**Figure 4.** Motor learning curves for real-time experiment. **(A)**-**(B)** Performance scores received by participants after trial end are plotted against the experimental block number for **(A)** EMG and **(B)** glove control; **(C)** comparison of early versus late blocks for grouped participants (able-bodied and amputee); **(D)** one-to-one comparisons of early versus late blocks averages for all participants. Points, means; error bars, 95% confidence intervals estimated via bootstrapping (1000 iterations); triple asterisk, $p < 0.001$.

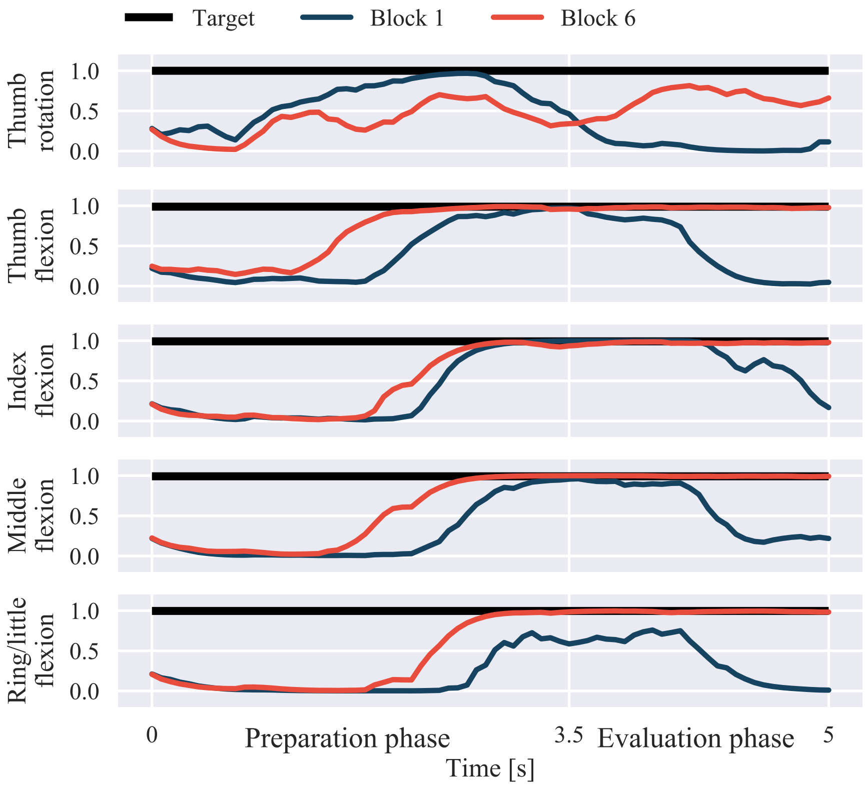


**Figure 5.** Example of learning effect on myoelectric control performance. Traces of target and actual digit positions of the robotic hand are shown for an able-bodied participant in blocks 1 and 6 of the experiment. The shown traces correspond to the full cylindrical grip (Supplementary Fig. 2N). The received performance scores at the end of the trials were 9.59% and 84.32% for blocks 1 and 6, respectively.

#### Supplementary Movies

Two movies are provided showing one amputee participant performing two blocks of the real-time posture matching task: Supplementary Movie S1, block 1; Supplementary Movie S2, block 6.

### Supplementary Methods

#### Mapping CyberGlove II measurements to degrees of actuation of the IH2 Azzurra hand

A linear mapping between the measurements of the 18-degree of freedom (DOF) CyberGlove II (CyberGlove Systems LLC) and the degrees of actuation (DOAs) of the IH2 Azzurra hand (Prensilia s.r.l.) was used. Due to cross-coupling between the data glove sensors[1,2], the mapping was identified empirically and its quality was subsequently verified during pilot experiments involving teleoperating the robotic hand in real-time using the data glove.

Let $x \in \mathbb{R}^{18}$ denote the calibrated measurements returned by the data glove, and $y \in \mathbb{R}^5$ denote the digit position vector of the DOAs of the hand. The elements in $y$ are ordered as follows: $y_1$, thumb rotation; $y_2$, thumb flexion; $y_3$, index flexion; $y_4$, middle flexion; $y_5$, ring/little flexion. The calibrated data glove measurements are then mapped onto robotic digit positions via a linear mapping:

$$y = Ax. \tag{1}$$

The transformation matrix $A$ was selected as follows:

$$A^\top = \begin{bmatrix} 0.639 & 0 & 0 & 0 & 0 \\ 0.383 & 0 & 0 & 0 & 0 \\ 0 & 1 & 0 & 0 & 0 \\ -0.639 & 0 & 0 & 0 & 0 \\ 0 & 0 & 0.4 & 0 & 0 \\ 0 & 0 & 0.6 & 0 & 0 \\ 0 & 0 & 0 & 0.4 & 0 \\ 0 & 0 & 0 & 0.6 & 0 \\ 0 & 0 & 0 & 0 & 0 \\ 0 & 0 & 0 & 0 & 0.1667 \\ 0 & 0 & 0 & 0 & 0.3333 \\ 0 & 0 & 0 & 0 & 0 \\ 0 & 0 & 0 & 0 & 0.1667 \\ 0 & 0 & 0 & 0 & 0.3333 \\ 0 & 0 & 0 & 0 & 0 \\ 0 & 0 & 0 & 0 & 0 \\ -0.19 & 0 & 0 & 0 & 0 \\ 0 & 0 & 0 & 0 & 0 \\ 0 & 0 & 0 & 0 & 0 \end{bmatrix} \tag{2}$$

#### References


1. Steffen, J., Maycock, J. & Ritter, H. Robust dataglove mapping for recording human hand postures. In *Proc. Int. Conf. Intell. Robots Appl.*, 34–45 (Springer, 2011).

2. Wang, Y. & Neff, M. Data-driven glove calibration for hand motion capture. In *Proc. ACM SIGGRAPH/Eurographics Symp. Comp. Anim.*, 15 (ACM Press, 2013).
